# Extracellular NAD(P) links hypersensitive response to localized acquired resistance

**DOI:** 10.64898/2026.01.13.699113

**Authors:** Fiona M. Harris, Cheng Liu, Qingcai Liu, Zhonglin Mou

## Abstract

Effector-triggered immunity (ETI) generates cell non-autonomous signals that activate localized acquired resistance (LAR) in neighboring cells and systemic acquired resistance (SAR) in distant tissues. While SAR signaling has been extensively characterized, the molecular basis of LAR remains poorly understood. Here, we identify extracellular nicotinamide adenine dinucleotide (eNAD(P)) as a key damage-associated molecular pattern (DAMP) that mediates LAR. Using genetic, physiological, and cell death assays, we show that eNAD(P) and its receptor complex restrict hypersensitive response (HR)-associated cell death, promote cell survival, and are required for avirulent pathogen-induced LAR. In contrast, LAR is independent of the major SAR mobile signals N-hydroxypipecolic acid and azelaic acid, as well as certain other characterized DAMP pathways. Notably, ETI-mediated resistance remains largely intact in eNAD(P) pathway mutants, indicating that ETI directly restricts pathogens while simultaneously generating eNAD(P) signals to activate LAR. Our findings establish LAR as a mechanistically distinct immune layer and link HR-associated damage to eNAD(P)-dependent local immune reinforcement.

## Main

Plant immunity at infection sites is mediated by two spatially and temporally distinct responses: pattern-triggered immunity (PTI) and effector-triggered immunity (ETI) (Jones and Dangl, 2006). PTI is activated by cell-surface pattern recognition receptors (PRRs) that detect conserved microbial signatures, known as pathogen-associated molecular patterns (PAMPs). PTI effectively prevents most potential pathogens from colonizing the plant. However, successful pathogens have evolved effector proteins that suppress PTI by targeting key immune signaling components, leading to effector-triggered susceptibility. To counteract this, plants have developed intracellular nucleotide-binding leucine-rich repeat (NLR) receptors that detect these effectors or their molecular consequences, activating effector-triggered immunity (ETI). ETI is a robust immune response that generates cell non-autonomous signals, which spread from the infection site to surrounding and distant cells. These signals activate localized acquired resistance (LAR), a strong but transient defense that restricts pathogen spread in cell layers adjacent to the site of infection (Ross, 1961), and systemic acquired resistance (SAR), a moderate yet long-lasting immune priming mechanism that enhances future defense responses in distal tissues (Fu and Dong, 2013).

A hallmark of strong ETI is the hypersensitive response (HR) that restricts local pathogen proliferation by producing antimicrobial compounds and depriving biotrophic pathogens of nutrients (Balint-Kurti, 2019). In the context of cell non-autonomous responses, LAR establishment and ETI-mediated disease resistance requires NLR activation but not necessarily the presence of macroscopic HR (Jacob et al., 2023). Similarly, pathogens that do not trigger a dominant ETI/HR response can still activate SAR, albeit with delayed kinetics, suggesting that HR is dispensable for both LAR and SAR but may serve to potentiate secondary immune signaling. These findings raise fundamental questions about the precise role of HR in ETI-induced resistance, as well as any mechanistic distinctions between LAR and SAR.

A recent perspective suggests that HR is not the goal but rather a consequence of strong ETI signaling, with its primary function being the release of cell non-autonomous signals that activate LAR (Jacob et al., 2023). In this model, LAR is proposed to be the primary driver of immunity during ETI, as pathogens fail to establish niches in the apoplast of adjacent cells following LAR activation. The underlying signals responsible for LAR establishment remain unclear. Although LAR-activated tissues accumulate salicylic acid (SA), SA alone is insufficient to trigger LAR (Costet et al., 1999). Importantly, the exact nature of the cell non-autonomous signal that induces LAR remains unclear, but has been proposed to be a damage-associated molecular pattern (DAMP) released during HR or as a result of membrane permeation by reactive oxygen species (ROS) (Jacob et al., 2023).

One such candidate DAMP is nicotinamide adenine dinucleotide (NAD) and its phosphorylated form, NADP. These universal electron carriers are present at millimolar concentrations inside cells, primarily functioning in metabolic reactions and intracellular signaling. During infection, pathogen-induced damage to plant cells causes NAD(P) to leak into the extracellular space. Once in the apoplast, extracellular NAD(P) (eNAD(P)) is detected by uninfected cells and initiates defense, particularly during SAR establishment (Li et al., 2023; Wang et al., 2019). As well, when infiltrated at physiological concentrations, NAD(P) strongly suppresses HR during avirulent pathogen challenge without compromising resistance (Zhang and Mou, 2012), suggesting that exogenously applied NAD(P) enhances immunity while preserving cell viability. We anticipate that cell survival is a tenant of LAR establishment, given that SA accumulation is associated with cell survival and that there is a biological need for local HR restriction during ETI responses. As eNAD(P) is a primary DAMP that is rapidly released into the apoplast during infection to induce SA-mediated resistance and cell survival, we hypothesized that it might be an LAR signaling molecule.

To confirm that NAD(P) functions in the apoplast to promote cell survival, we treated the *fin4-3* mutant (Macho et al., 2012), defective in the first irreversible step of *de novo* NAD biosynthesis, with either 4 mM nicotinic acid (NA), a cell-permeable NAD precursor that is converted intracellularly via the salvage pathway, or 1 mM NAD^+^, which is membrane impermeable. NAD^+^, but not NA, significantly reduced *Pseudomonas syringae* pv. *tomato* DC3000 (*Pst*) carrying the effector gene *HopZ1a* (*Pst avrHopZ1a*)- and *Pst avrRps4*-induced ion leakage (an indicator of loss of membrane integrity and cell death) in wild-type (WT) and *35S:FIN4/fin4-3* plants (Figures 1A and 1B). Furthermore, while NA restored *Pst avrHopZ1a*- and *Pst avrRps4*-induced ion leakage to wild-type levels in *fin4-3*, it failed to enhance cell survival significantly, unlike NAD^+^ treatment (Figures 1A and 1B). These findings indicate that NAD(P) promotes cell survival specifically when present in the apoplast.

**Figure 1.**
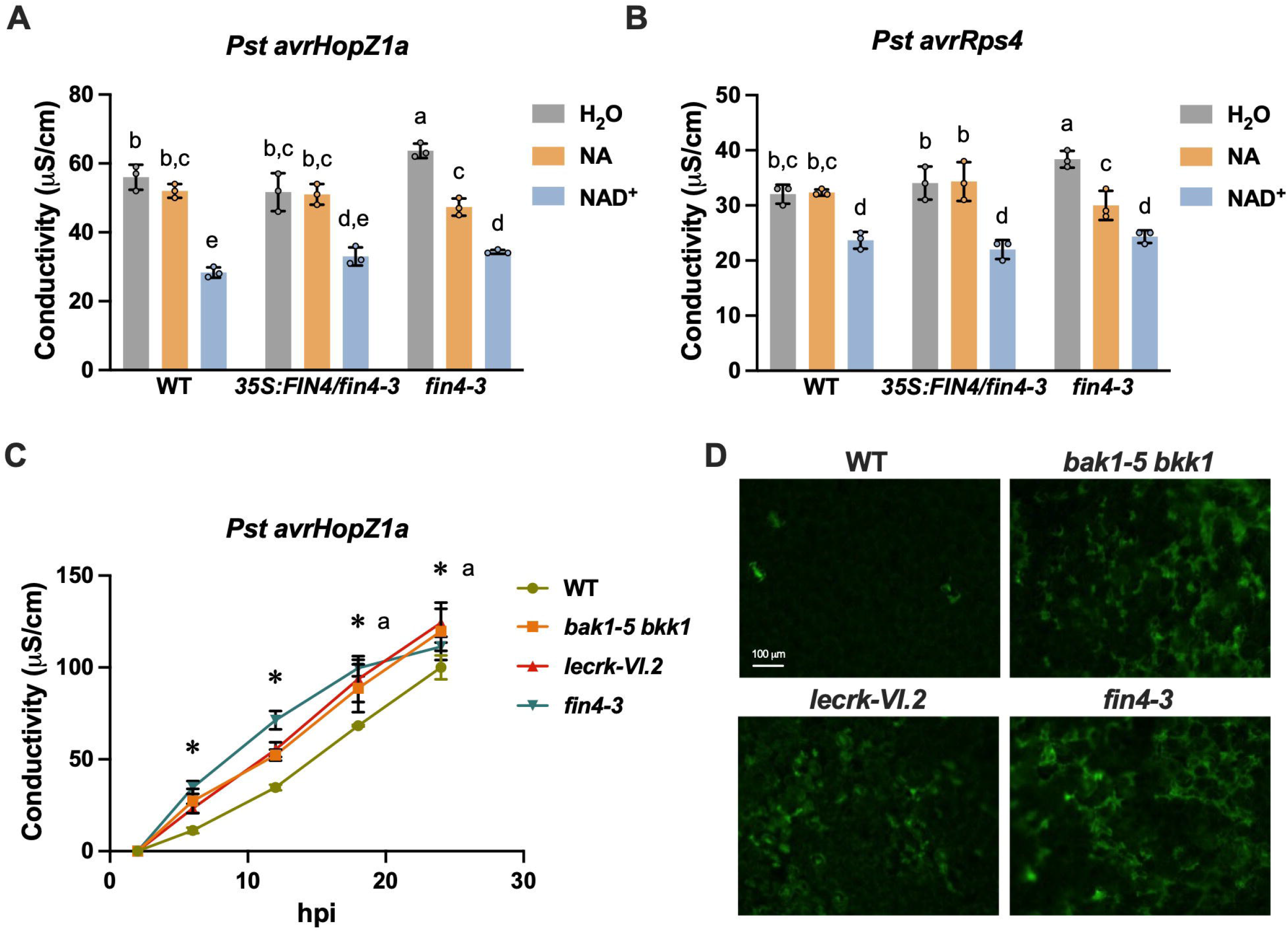
Endogenous eNAD(P) restricts HR-mediated cell death. **A and B)** NA- and NAD^+^-induced suppression of *Pst avrHopZ1a*-(**A)** and *Pst avrRps4*-triggerd (**B**) ion leakage. Leaves of 4-week-old wild-type (WT), *35S:FIN4/fin4-3*, and *fin4-3* plants were infiltrated with ddH_2_O, 4 mM NA, or 1 mM NAD^+^. Four hr later, the infiltrated leaves were inoculated with *Pst avrHopZ1a* or *Pst avrRps4* (OD_600_ = 0.1). Ion leakage (conductivity) was measured at 18 hr post-inoculation (hpi). Bars represent means ± standard deviation (SD) (n = 3). Different letters denote significant differences (*P* < 0.05; one-way ANOVA). **C)** Ion leakage from leaves of the indicated genotypes inoculated with *Pst avrHopZ1a* (OD_600_ = 0.025). Conductivity was measured at 2, 6, 12, 18, and 24 hpi. Error bars represent SD (n = 3). Asterisks denote significant differences (\**P* < 0.05; Student’s t-test). a, ion leakage of the *bak1-5 bkk1* mutant was not significantly different from that of the wild type at 18 and 24 hpi. **D)** Autofluorescence imaging of *Pst avrHopZ1a*-triggered HR-mediated cell death in the indicated genotypes. Leaves of 4-week-old plants were infiltrated with *Pst avrHopZ1a* (OD_600_ = 0.025). Autofluorescence images (488 nm excitation and 507 nm emission) were captured from the abaxial side at 24 hpi using 20× magnification. Green fluorescence indicates dying or dead cells.

We next investigated avirulent pathogen-induced cell death in *fin4-*3, which accumulates reduced eNAD(P) levels after infection (Li et al., 2023), as well as the eNAD(P) receptor complex mutants, *lecrk-VI*.*2* and *bak1-5 bkk1* (Schwessinger et al., 2011; Wang et al., 2019). All three mutants exhibited significantly higher ion leakage than the wild type upon infection with *Pst avrHopZ1a* (Figure 1C), suggesting that eNAD(P) and its receptor complex regulate HR-associated cell death. To further investigate the altered cell death phenotype, we assessed whether autofluorescence imaging using a standard GFP filter (488 nm excitation, 507 nm emission) could serve as a reliable tool for detecting single-cell HR, as the accumulation and release of secondary metabolites during cell death leads to autofluorescence in the blue-green spectrum (Koga et al., 1988; Donaldson, 2020). We imaged cell death at 24 h post-inoculation (hpi) in wild type and *zar1-3*, a mutant of the resistance protein recognizing HopZ1a, and detected autofluorescence exclusively in the wild type under *Pst avrHopZ1a* treatment, not *Pst* harboring the empty vector (Supplemental Figure 1), confirming that this approach successfully visualizes HR at the cellular level.

Notably, *Pst avrHopZ1a*-triggered autofluorescence patterns differed markedly between genotypes of interest (Figure 1D). In wild type, autofluorescence was confined to discrete, single-cell-sized regions with well-defined boundaries, consistent with controlled HR. In contrast, *lecrk-VI*.*2, bak1-5 bkk1*, and particularly *fin4-3* exhibited diffuse and expanded autofluorescence, indicative of uncontrolled cell death. These findings align with ion leakage results, reinforcing that eNAD(P) and its receptor complex restrict HR-associated cell death.

To assess the role of eNAD(P) in LAR, we adapted Ross’s classic LAR assay for *Arabidopsis* (Ross, 1961). Leaves were first inoculated with low concentrations of avirulent bacterial pathogens (*Pst avrHopZ1a, Pst avrRps4*, and *Pst avrRpm1*) and subsequently challenged 4 h later with the virulent *P. syringae* pv. *maculicola* ES4326 strain carrying a *luxCDABE* cassette (*Psm_lux*). The *Pst* strain harboring pUCP20 (empty vector) was included as a control. Since host plant cells undergoing HR are typically separated by multiple cell layers following low-dose infiltration of avirulent *Pst* (Rufian et al., 2018), virulent pathogens can still colonize the apoplast around these cells. To validate this methodology, we tested the LAR phenotype of the SA pathway mutants *sid2* and *npr1-3*, defective in SA biosynthesis and signaling, respectively. *Pst avrHopZ1a* and *Pst avrRps4*, but not *Pst*, induced strong LAR in the wild type, while *sid2* and *npr1-3* failed to mount an effective LAR response (Figures 2A and 2B), supporting a critical role for SA in LAR (Costet et al., 1999). Notably, LAR induced by *Pst avrHopZ1a, Pst avrRps4*, and *Pst avrRpm1* was significantly compromised in the eNAD(P) pathway mutants (*fin4-3, lecrk-VI*.*2*, and *bak1-5 bkk1*) (Figures 2C-2E), indicating that eNAD(P) is a critical signaling molecule in LAR. Consistent with this conclusion, *Pst avrHopZ1a*-induced expression of *PR1*, a marker gene of SA signaling, was impaired in these mutants (Supplemental Figure 2). However, resistance to the three avirulent pathogens remained largely intact in these mutants (Supplemental Figures 3A-3C), supporting the idea that ETI confers direct resistance while simultaneously generating signals for LAR activation.

**Figure 2.**
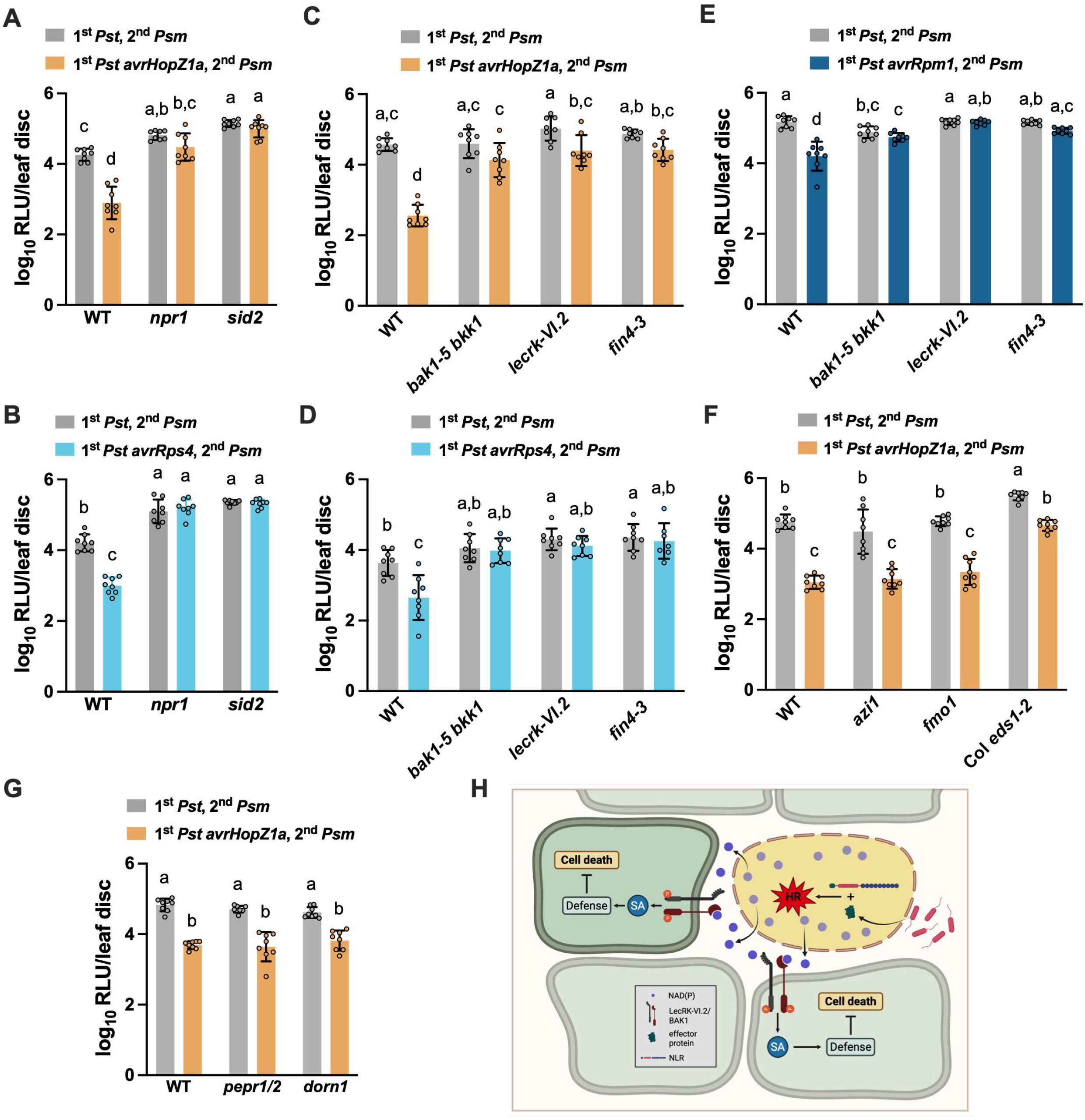
eNAD(P) and its receptor complex mediate localized acquired resistance. **A to G)** Leaves of 4-week-old wild-type (WT) and the indicated *Arabidopsis* mutant plants were first (1^st^) infiltrated with *Pst* or *Pst* harboring *avrHopZ1a, avrRps4*, or *avrRpm1* (OD_600_ = 0.0005), Four hr later, the infiltrated leaves were challenge-inoculated (2^nd^) with *Psm* (*Psm_lux*, OD_600_ = 0.001). *Psm* populations were measured at 2.5 d post-inoculation (dpi). Bars represent means ± SD (n = 8). Different letters denote significant differences (*P* < 0.05; one-way ANOVA). RLU, relative light unit. **H)** Proposed model illustrating how eNAD(P) mediates LAR. Recognition of pathogen-delivered effector proteins or their activities triggers ETI and the associated HR. HR cell death leads to the release of NAD(P) into the apoplast, where it is perceived by the cell-surface LecRK-VI.2/BAK1 receptor complexes of neighboring cells. This perception activates SA-dependent defense signaling, which suppresses further cell death and restricts pathogen spread in surrounding tissues.

Given that eNAD(P) functions as an SAR signaling molecule, we hypothesized that other major SAR mobile signals might also contribute to LAR. However, *Pst avrHopZ1a*-induced LAR remained intact in *fmo1* and *azi1* mutants (Figure 2F), which are defective in the biosynthesis and signaling of the SAR mobile signals N-hydroxypipecolic acid and azelaic acid, respectively. We also investigated potential roles of other characterized DAMPs in LAR signaling. Interestingly, *Pst avrHopZ1a*-induced LAR is unaffected in *dorn1* and *pepr1/2*, the receptor mutants of extracellular ATP and AtPeps, respectively (Figure 2G). These findings indicate that LAR is mechanistically distinct from SAR and that not all DAMPs contribute to LAR.

SAR is thought to result from the synergistic action of multiple mobile signals produced at the primary infection site (Kachroo and Kachroo, 2020). However, the concentration of any single signal in systemic tissues is likely too low to activate SAR independently. A signal amplification loop, in which each signal reinforces the production of others, has been proposed as a key mechanism in SAR (Kachroo and Kachroo, 2020). This amplification regulates ROS accumulation, enabling ROS to permeate the plasma membrane without inducing excessive cell death, thereby facilitating controlled NAD(P) leakage (Li et al., 2023). The resulting eNAD(P) then acts through its receptor complex to establish SAR.

In contrast, during ETI in response to avirulent pathogens, a rapid and intense ROS burst exceeds the lethal threshold, leading to both eNAD(P) accumulation and localized cell death.

However, this ROS burst is not entirely cell-autonomous and can diffuse into neighboring cells to facilitate NAD(P) release. Since this response is robust, transient, and spatially restricted, LAR is characterized by its strong yet short-lived nature.

The role of HR-associated cell death in ETI remains to be fully understood (Jacob et al., 2023). Genetic evidence suggests that HR is not strictly required for resistance against biotrophic and hemibiotrophic pathogens (Jacob et al., 2023), raising the question of why HR has been evolutionarily conserved in ETI. By linking HR, eNAD(P), and LAR, our findings support the idea that, beyond pathogen restriction, HR potentiates LAR signaling by enhancing eNAD(P) accumulation (Figure 2H). This study reconciles the apparent dispensability of HR in resistance with its evolutionary conservation. Future research into NLR signaling, ROS dynamics, and eNAD(P) signaling will be crucial for developing genetic modifications and breeding strategies to enhance disease resistance in crops.

## Author contributions

F.M.H. and Z.M. conceived the project. F.M.H., C.L., and Q.L. performed the experiments and analyzed the data. F.M.H. and Z.M. prepared figures and wrote the manuscript. All authors provided comments for the manuscript before submission.

## Acknowledgements

We are grateful to Dr. Darrell Desveaux (University of Toronto, Canada) for sharing *P. syringae* pv. *tomato* DC3000 strains carrying the empty vector pUCP20 and hopZ1apro:hopZ1a-HA in pUCP20, Dr. Cyril Zipfel (University of Zurich, Switzerland) for *bak1-5 bkk1* seeds, Dr. Jean Greenberg (University of Chicago, USA) for *azi1* seeds, and Dr. Yusuke Saijo (Nara Institute of Science and Technology, Japan) for *pepr1 pepr2* seeds. The T-DNA insertion lines SAIL_1145_B10 (*fin4-3*), SALK_070801 (*lecrk-VI*.*2-1*), and SALK_042209 (*dorn1-3*) were ordered from the Arabidopsis Biological Resource Center.

