## Supplemental figures and methods for "Extracellular NAD(P) links hypersensitive response to localized acquired resistance"

**Supplemental figure and figure legends**

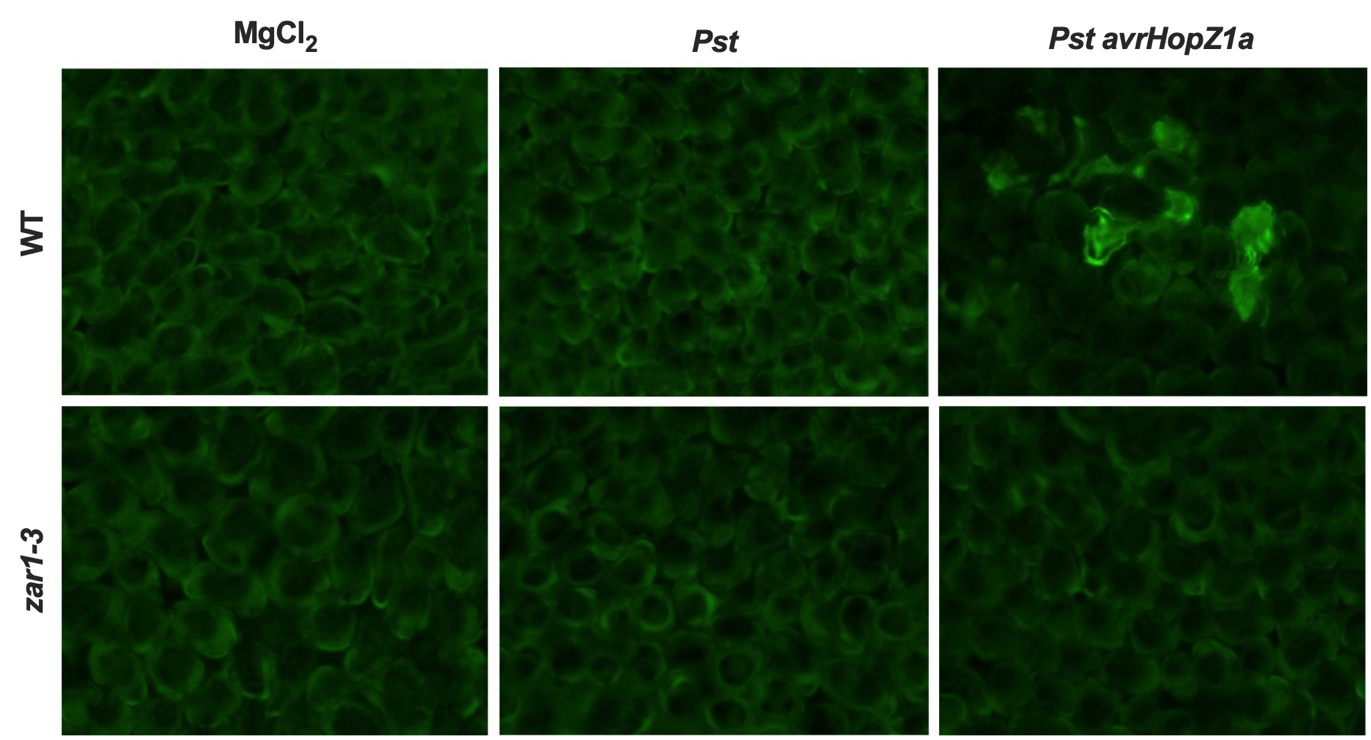

**Supplemental Figure 1 Validation of blue light imaging of autofluorescence during HR-mediated cell death.** Leaves of 4-week-old wild-type (WT) and *zar1-3* mutant plants were infiltrated with *Pst avrHopZ1a* (OD_600_ = 0.025). Autofluorescence from dying cells was imaged at 24 hpi using the GFP channel (488 nm excitation and 507 nm emission) of a Zeiss Axios Observer fluorescence microscope at 20× magnification. Image contrast was enhanced using autothresholding.

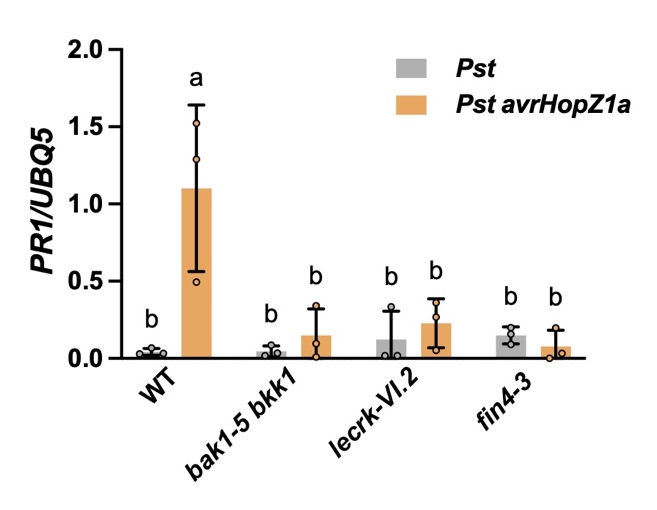

**Supplemental Figure 2 *Pst avrHopZ1a*-induced *PR1* expression in eNAD(P) pathway mutants.** Leaves of 4-week-old wild-type (WT), *lecrk-VI.2*, *bak1-5 bkk1*, and *fin4-3* mutant plants were infiltrated with a bacterial suspension (OD_600_ = 0.0005) of *Pst* or *Pst avrHopZ1a.* Leaf tissues were collected 4 h later for qPCR analysis. Expression levels were normalized against the constitutively expressed *UBQ5*. Bars represent means ± SD (n = 3). Different letters denote significant differences (p < 0.05; one-way ANOVA).

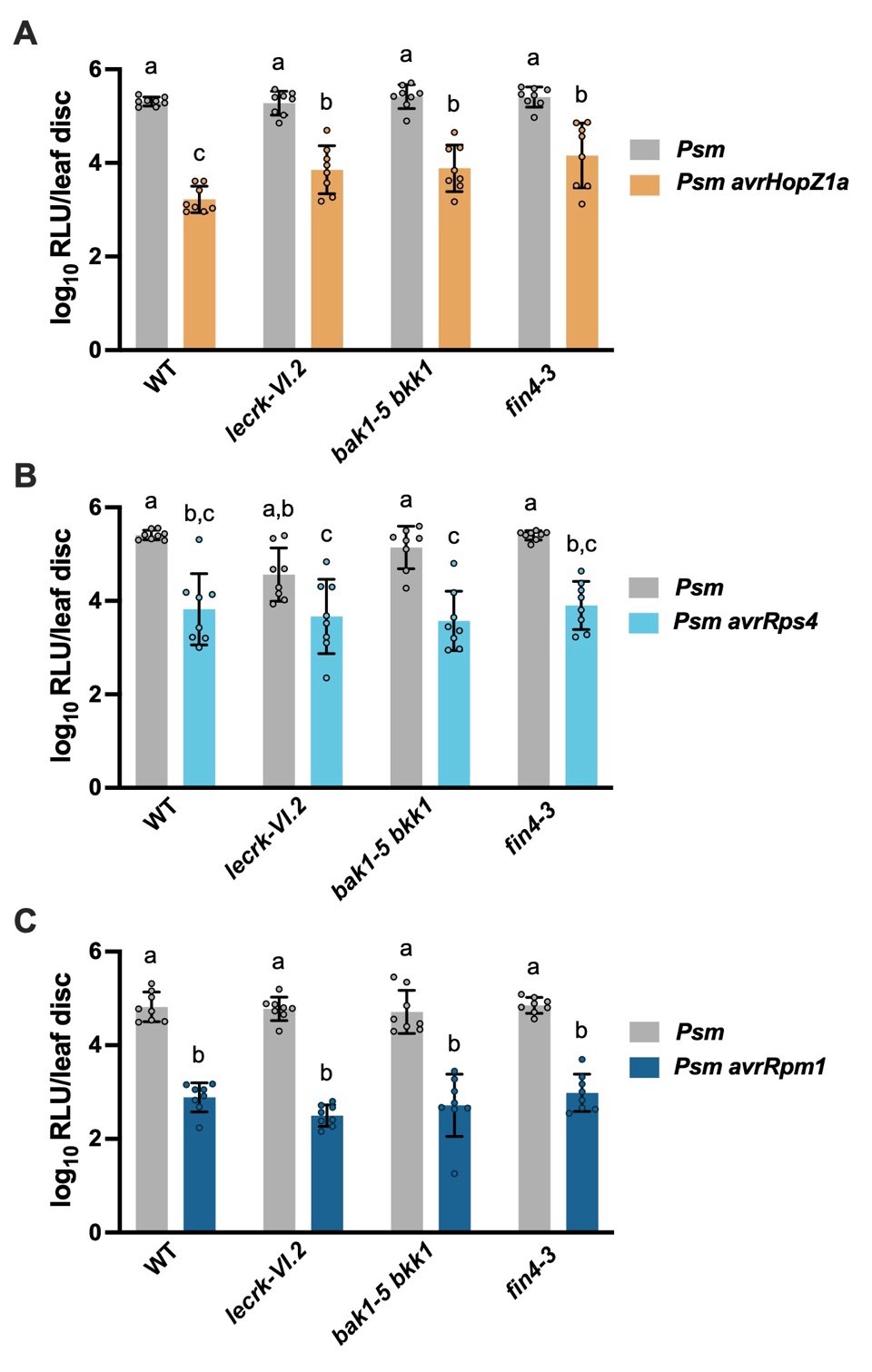

**Supplemental Figure 3 eNAD(P) and its receptor complex are dispensable for ETI-mediated direct pathogen restriction. (A**-**C)** Leaves of 4-week-old wild-type (WT), *lecrk-VI.2*, *bak1-5 bkk1*, and *fin4-3* mutant plants were infiltrated with *Psm*, *Psm avrHopZ1a* (**A**), *Psm* *avrRps4* (**B**), or *Psm* *avrRpm1* (**C**) (OD600 = 0.001). Bacterial populations were measured at 2.5 dpi. Bars represent means ± SD (n = 8). Different letters denote significant differences (*P* < 0.05; one-way ANOVA). RLU, relative light unit.

**Supplemental methods**

**Plant materials and growth conditions**

*Arabidopsis* plants were grown in soil for 4 weeks under greenhouse conditions (24/22°C day/night temperatures, ~60% relative humidity, and a 14/10 h photoperiod). All mutant lines are in Columbia (Col-0) background and have been previously used (Li *et al.*, 2023).

**Transformation of *Psm_lux* strains**

The previously published *Psm_lux* strain (Li *et al.*, 2023) was streaked onto King’s B media containing 50 μg/mL streptomycin and grown at 28°C for 3 d. A single colony was used to inoculate 6 mL of liquid King’s B medium supplemented with streptomycin (50 μg/mL) in a 50 mL tube with a loose-fitting cap and cultured overnight at 28°C with shaking. The culture was pelleted by centrifugation at 4,000 rpm for 5 min, washed with 5 mL room temperature sterile water, pelleted again, and resuspended in ice-cold 300 mM sucrose prior to flash-freezing in liquid nitrogen. Fresh competent cells were electroporated with approximately 500 ng of pVSP61, pVSP61-avrRps4, pUCP20, or pUCP20-avrHopZ1a using a Gene Pulser Xcell PC system (Bio-Rad) with the parameters: 5 kV, 25 μF, and 200 Ω, and then recovered in 1 mL King’s B for 4 h prior to plating on King’s B containing streptomycin and 50 μg/mL kanamycin.

**Conductivity measurement**

For local tissues, two fully expanded leaves of 4-week-old *Arabidopsis* plants were infiltrated on the abaxial side with 1 mM NAD^+^, pH 5.7, or mock (H_2_O) with a 1 mL needleless syringe 4 h prior to inoculation with a bacterial suspension (OD_600_ = 0.2) of avirulent *Pst* and *Psm_lux* strains carrying either *avrRps4* or *avrHopZ1a*. Twelve leaf discs per genotype/treatment were sampled approximately 20 min post pathogen inoculation and washed 3 times in 25 mL sterile ddH_2_O for 30 min. Four leaf discs were placed into 5 mL sterile ddH_2_O, and 50 uL was taken for conductivity measurement every 6 h over 24 h using a HORIBA conductivity meter (model EC-22).

**Autofluorescence measurement of cell death**

Leaves of 4-week-old *Arabidopsis* plants were infiltrated with a *Pst* *avrHopZ1a* suspension (OD_600_ = 0.025) and sampled at 24 hpi. Leaf sections (~5×5 mm) were excised using surgical scissors, avoiding veins, and placed abaxial side up under coverslips with 250 uL ddH_2_O. Samples were imaged on a Zeiss Axios Observer 3, using the GFP channel (488 nm excitation and 507 nm emission) at 20 × magnification. Appropriate Z-stack limits were established, and auto-thresholding was used to optimize image resolution. Four images were taken at unique, resolvable (flat) areas per excision, and three biologic replicates were sampled per treatment per genotype.

**Quantification of LAR and ETI**

Leaves of 4-week-old *Arabidopsis* plants were infiltrated with a bacterial suspension (OD_600_ = 0.0005) of the avirulent strain *Pst* *avrHopZ1a*, *Pst avrRPS4*, or *Pst avrRpm1*. For *Pst* *avrHopZ1a* and *Pst avrRPS4*, the infiltrated leaves were challenged 4 h later with *Psm_lux* (OD_600_ = 0.001)*.* As *Pst avrRpm1-*triggered HR occurs earlier than *Pst* *avrHopZ1a* and *Pst avrRPS4*, *Pst avrRpm1*-infiltrated leaves were challenged 2 h later. Eight leaf discs were sampled per treatment per genotype at 2-2.5 d post-challenge inoculation and placed in a white 96-well plate containing 150 uL ddH_2_O. Luminescence was measured using a GloMax Discover Microplate Reader (Promega).

For ETI assay, leaves of 4-week-old *Arabidopsis* plants were infiltrated with a bacterial suspension (OD_600_ = 0.0005) of the virulent strain *Psm_lux* or avirulent *Psm_lux* strains carrying *avrHopZ1a*, *avrRps4*, or *avrRpm1* using a needleless syringe. Eight leaf discs were sampled per treatment per genotype at 2-2.5 d post-infiltration and placed in a white 96-well plate containing 150 uL ddH_2_O. Luminescence was measured as above.

**RNA extraction and qPCR analysis**

Leaves of 4-week-old *Arabidopsis* plants were infiltrated with a bacterial suspension (OD_600_ = 0.0005) of *Pst* or *Pst avrHopZ1a.* Four h later, four 50 mg samples, each from an independent plant per treatment, were collected, weighed, and flash frozen. RNA was extracted using the E.Z.N.A. Plant RNA Kit (Omega Bio-Tek, R6827-01) per the manufacturer’s instructions. Briefly, frozen tissues were homogenized via bead beating at 1500 rpm for 30 s, then thawed and vortexed at room temperature in 500 uL RB Buffer containing 2% β-mercaptoethanol. Resuspended tissue was cleared by centrifugation in a Homogenizer Mini Column (14,000 × g, 5 min, room temperature (R.T.)), and eluant was vortexed with 1:1 (v:v) with 70% ethanol. Sample was run through the HiBind RNA Mini Column (12,000 × g, 1 min, R.T.) until consumed and then washed on the column with 500 uL RNA Wash Buffer I, followed by 500 uL RNA Wash Buffer II (10,000 × g, 30 s, R.T.). The column was dried via centrifugation at 15,000 rpm for 2 min, then the RNA was eluted in 50 uL nuclease-free water.

Reverse transcription (RT) was performed using the LunaScript RT SuperMix Kit (NEB, E3010), per the user instructions. qPCR reactions were performed using 2 uL of cDNA from above and the AB Clonal 2× Universal SYBR Green Fast qPCR Mix (RK21203) on a QuantStudio 3 Real-Time PCR system (Applied Biosystems), per the manufacturer’s instructions. Expression levels were determined using the 2^-ΔCt^ method relative to *UBQ5* as an internal control. The primers used for qPCR were reported previously (Wang *et al.*, 2019).
